# TP-MAP - an Integrated Software Package for the Analysis of 1D and 2D Thermal Profiling Data

**DOI:** 10.1101/2021.02.22.432361

**Authors:** Felix Feyertag, Kilian V.M. Huber

## Abstract

Thermal profiling (TP) has emerged as a promising experimental methodology for elucidating the molecular targets of drugs and metabolites on a proteome-wide scale. Here, we present the Thermal Profiling Meltome Analysis Program (TP-MAP) software package for the analysis and ranking of 1D and 2D thermal profiling datasets. TP-MAP provides a user-friendly interface to quickly identify hit candidates and further explore targets of interest via intersection and crosslinking to public databases.

## Introduction

Assessing molecular perturbations in living cells caused by drug treatment, environmental changes, genetic mutations or alterations in metabolic flux in an unbiased and proteome-wide manner constitutes a key challenge in chemical biology and systems pharmacology. Recently, thermal profiling (TP) has emerged as a novel approach for the unbiased interrogation of drug and metabolite effects in intact cells or cell lysate^1–3^. The underlying premise is built upon the notion that proteins tend to exhibit changes in thermal stability in ligand-bound versus unbound state. The cellular thermal shift assay (CETSA) was developed to elucidate drug-target interactions in living cells by heating compound treated and untreated cells and quantifying the change in aggregation or ‘melting’ behaviour between the two conditions for a given protein^4^. When applying multiple concentrations, isothermal dose response fingerprints (ITDRF) can be utilised to determine the dose-dependent response to a ligand and estimate binding affinities. TP combines the CETSA and ITDRF approaches with quantitative multiplexed mass spectrometry to enable the large-scale quantification of thousands of proteins, thereby significantly expanding the coverage of CETSA/ITDR from a single cognate target to the entire proteome in theory. Over the last couple of years, several studies have successfully utilised 1-dimensional (1D) TP (i.e. one fixed concentration of ligand versus temperature) and 2-dimensional (2D) TP (i.e. dose-response matrix with multiple concentrations and temperatures) for various purposes. TP was first applied for the identification of global target engagement of drugs^1,3–7^, and has since also been extended to areas such as investigating drug-induced re-wiring of protein interaction networks^8–11^, the identification of protein complex subunits^12^, and the characterisation of metabolic processes in humans^13,14^ and bacteria^15,16^.

Bioinformatic workflows for analysing 1D TP commonly involve fitting sigmoid-like melting curves to treatment and vehicle replicates, and measuring the thermal shift observed between the two^5,15^; more recently, non-parametric methods have been reported^17^. For 2D TP, concentration-dependent dose-response curves are generally fitted for each temperature^6,14^, and recent studies have extended the use of non-parametric methods from 1D thermal profiling to the analysis of 2D datasets^17,18^.

In order to further facilitate the analysis of TP datasets and adoption of the TP method in general, we developed the Thermal Profiling Meltome Analysis Program (TP-MAP) which features a user-friendly graphical user interface for analysing 1D and 2D TP datasets. TP-MAP offers functionality to import protein abundance data, calculates ratios and normalises the data, and provides the user with a summary table that can be inspected to identify proteins of interest, including ranking proteins by apparent stabilisation. For 1D datasets, ranking is based on thermal shift and curve fit quality, whereas for 2D datasets we present a new approach for ranking based on a hill-climbing algorithm. In addition, TP-MAP supports downstream analysis options including exporting proteins of interest to external databases for further exploration of e.g. protein-protein interaction networks and protein complexes.

## Results

### 1D Thermal Profiling Analysis

For 1D thermal profiling, TP-MAP fits a denaturation curve and determines the temperature at which 50% denaturation is observed (T_M_). In addition to T_M_, the 1D thermal profiling score considers the quality of the curve fit and reproducibility metrics. As the scoring may require adjusting for different datasets due to differences in data quality, experimental setup and target proteome, the score weighting can be adjusted interactively to prioritise T_M_ shift or the quality of curve fit (see methods). The default weighting is set at 70% and can be shifted higher to prioritise potential targets with large T_M_ shifts or lower for targets with small but reproducible T_M_ shifts.

To evaluate TP-MAP’s ability to correctly identify hits from 1D TP experiments, we selected previously published datasets for intact K562 cells treated with either two different concentrations of the clinical kinase inhibitor dasatinib (0.5 μM and 5 μM) versus DMSO surveyed over 10 temperature points^3^. As a reference for true hits we extracted and mapped known dasatinib drug targets^19^.

First, we assessed the outcome of shifting the weight of the TP-MAP score on target identification. Using a 50% T_M_ shift weighting, TP-MAP successfully identified five known dasatinib kinase targets in the 0.5 μM dataset^19^ (BTK, ABL2, YES1, MAPK14 and MAPKAPK2), only two of which had previously been reported from thermal profiling experiments. Similarly, in the 5 μM dasatinib dataset, five kinase targets were picked up (MAPK14, BTK, CSK, GAK and YES1) when the TP-MAP weighting was set to 70% (Supplementary Figure 1). Interestingly, BTK and YES1 were found to have a negative T_M_ shift in both 0.5 μM and 5 μM datasets. The high affinity dasatinib target BTK^20^ ranked highly when the weighting was low to prioritise curve fit over T_M_, ranking first in the 5 μM dataset when the weighting was set between 10-30% and second in the 0.5 μM dataset between 20-60%. Conversely, mitogen activated protein kinase MAPK14 (p38α) ranked first in the 5 μM dataset when setting the weight between 40-90% (Figure 1). Using the TP-MAP built-in functionality to explore PPI networks of identified targets via STRING^21^, indirect targets of dasatinib were revealed such as CRKL, INPPL1^19^ and STAT5B^22^, which also ranked highly using TP-MAP (Supplementary Figure 2).

**Figure 1.**
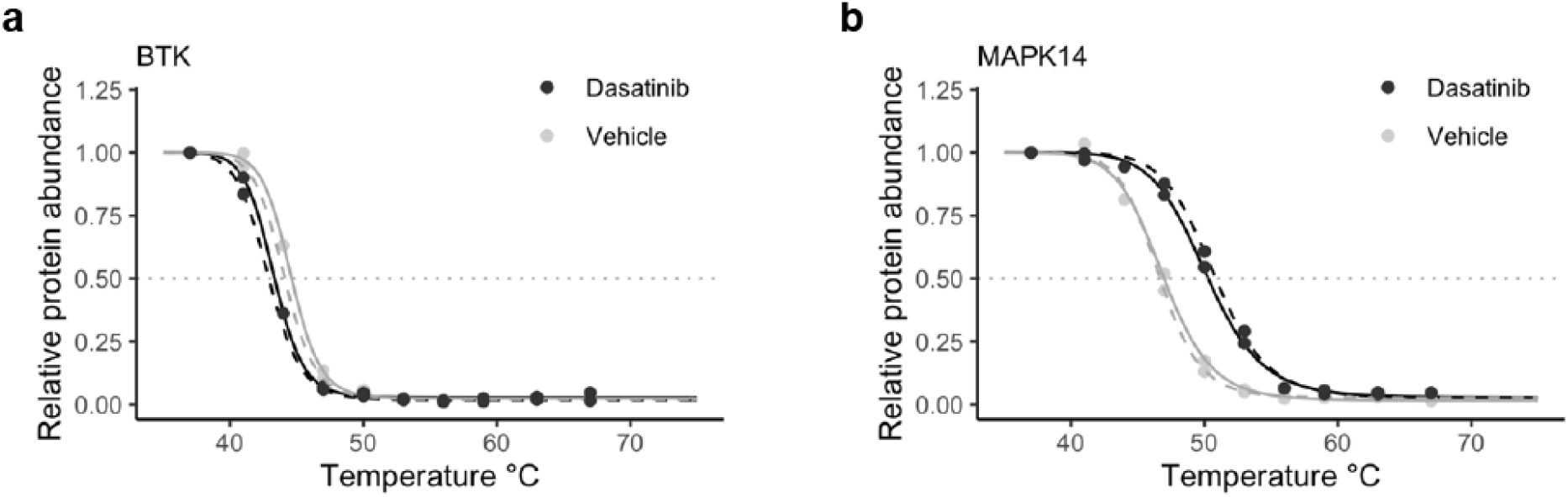
Melting curves for BTK (a) and MAPK14 (a) in intact cells treated with dasatinib 5 μM. BTK had a small but reproducible negative T_M_ shift and ranked first with a 1D weighting of 10-30% whereas MAPK14 ranked first when the weighting was shifted to prioritise T_M_ shift between 40-90%.

We further analysed a 1D thermal profiling dataset for the HDAC inhibitor panobinostat^5^ using TP-MAP. Out of 9 proteins known to have a thermal shift within this dataset, including six histone deacetylases (HDAC1, −2, −6, −7, −8, and −10), STX4, TTC38 and ZFYVE28 were ranked in the top 0.6% with weighting adjusted to 90% (Table 1).

**Table 1.**
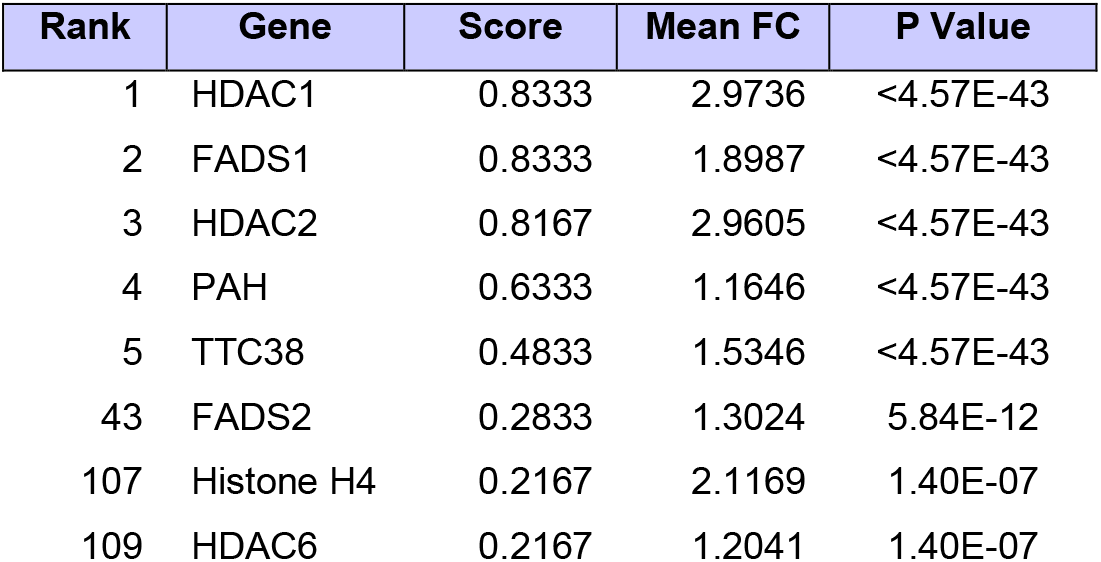
Ranking of targets and off-targets as well as indirectly affected proteins of panobinostat in intact HepG2 cells. Five of the direct targets appear as the top 5 stabilised proteins when ranked using the TP-MAP score (combined) and mechanistically-related indirect targets, e.g. histone H4, appear ranked in the top 2% of 8280 proteins.

| Rank | Gene | Score | Mean FC | P Value |
| --- | --- | --- | --- | --- |
| 1 | HDAC1 | 0.8333 | 2.9736 | <4.57E-43 |
| 2 | FADS1 | 0.8333 | 1.8987 | <4.57E-43 |
| 3 | HDAC2 | 0.8167 | 2.9605 | <4.57E-43 |
| 4 | PAH | 0.6333 | 1.1646 | <4.57E-43 |
| 5 | TTC38 | 0.4833 | 1.5346 | <4.57E-43 |
| 43 | FADS2 | 0.2833 | 1.3024 | 5.84E-12 |
| 107 | Histone H4 | 0.2167 | 2.1169 | 1.40E-07 |
| 109 | HDAC6 | 0.2167 | 1.2041 | 1.40E-07 |

We next investigated the ability of TP-MAP to classify proteins on a quantitative scale by investigating protein stabilisation among ATP-binding proteins using a dataset treated with Na-ATP^14^. We used a 1D TP dataset using Jurkat crude lysate treated with 2 mM Na-ATP heated between 37-67 °C in two replicates. We downloaded proteins from UniProt that were annotated with the gene-ontology term ATP-binding (GO:0005524), which resulted in 475 proteins being annotated as ATP-binding (12.69%). We evaluated the predictive power of TP-MAP in identifying ATP-binding proteins from the TP-MAP 1D score by means of varying percentage weights using receiver operator characteristic (ROC) curves. An optimal area-under-curve (AUC) was obtained using a weight of 70% with an AUC of 0.739 compared to an AUC of 0.722 when the weighting was set to 100% (Figure 2). A higher AUC of 0.771 was obtained using T_M_ shift when ranked from positive (stabilising) to negative (destabilising). This may be a consequence of the TP-MAP score using the absolute T_M_ shift, thereby not favouring stabilised proteins *a priori*. Indeed, if we limit our analysis to proteins with a positive mean T_M_ shift between replicates, AUC is increased using the 1D TP-MAP score at a weight of 85% to 0.799, which is an improvement compared to 0.789 using mean T_M_ shift alone.

**Figure 2.**
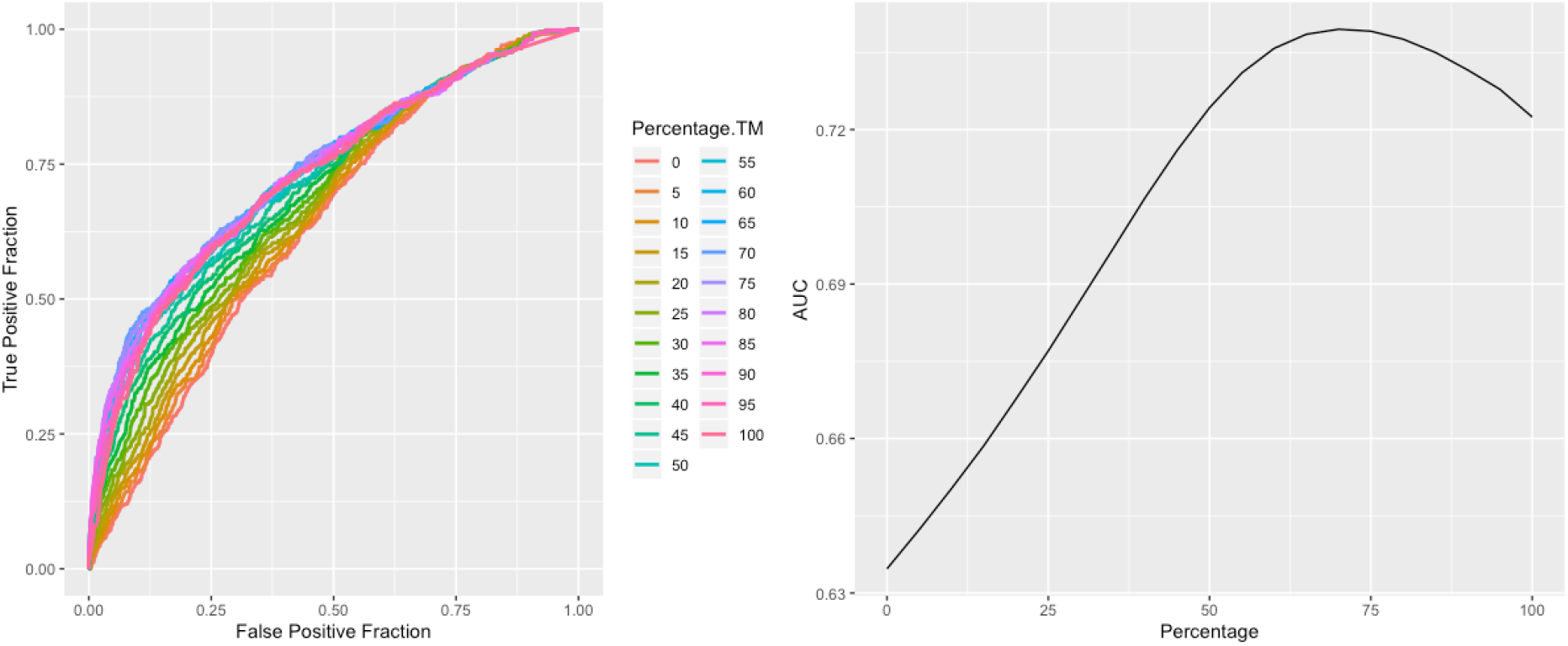
AUC curve classifying ATP-binding proteins in 1D thermal profiling datasets using the TP-MAP 1D score for weighting between 0 and 100%, where 0% only considers curve fit quality and 100% is equivalent to T_M_ shift. The optimal AUC for the ATP dataset is at 70%.

Taken together, these data suggest that a default weighting of 70% appears a reasonable starting point but should be adjusted as needed depending on the dataset.

### 2D Thermal Profiling Analysis

A challenge with 1D experiments is that often a multitude of proteins exhibit apparent substantial shifts which makes prioritisation of potential hits for follow-up experiments difficult even when using optimised scoring metrics. In order to address this issue and to facilitate the identification of true and relevant targets, 2D thermal profiling adds concentration as an additional dimension which allows for dose-dependent prioritisation of hits. However, most current methods for analysing 2D TP data do not provide an automated ranking for 2D datasets. TP-MAP incorporates a novel metric for scoring 2D TP datasets based on the assumption that concentration-dependent stabilisation or destabilisation will occur at an optimum temperature and concentration combination, and that adjacent temperature and concentration combinations will ascend or descend towards an optimum point (see methods). A percentile threshold can be set to determine the maximum and minimum fold change required for a peak or trough to be considered stabilising or destabilising, respectively. Analogously to the 1D datasets, we benchmarked a number of 2D TP datasets and found a percentile threshold of 80% to be a good default setting.

First, we used TP-MAP to analyse a previously published 2D TP dataset investigating the HDAC inhibitor panobinostat^6^ in HepG2 intact cells and cell lysate. Processing of the data with TP-MAP revealed 899 out of 8280 proteins that exhibited drug-induced stabilisation in intact cells (p-value < 0.05), and 517 out of 6971 proteins in cell lysate. Notably, the TP-MAP scoring method ranked the primary cognate targets HDAC1 and HDAC2 highly, with HDAC1 coming out first in both intact cells (score 0.83, p-value < 5.57 × 10^−43^) and cell lysate (score 0.83, p-value < 2.22 × 10^−16^), and HDAC2 ranked third in intact cells (score 0.82, p-value < 5.57 × 10^−43^) and second in cell lysate (score 0.67, p-value < 2.22 × 10^−16^). Other off-targets and mechanistically related indirect targets also ranked highly using TP-MAP (Tables 1 and 2).

**Table 2.** Ranking of targets and off-targets as well as indirectly affected proteins of panobinostat in HepG2 cell lysate. All six identified direct targets appear among the top 7 results when ranked using TP-MAP score (combined). As would be expected in cell lysate, indirect targets FADS2 and histone H4 do not change significantly in thermal stability.

| Rank | Gene | Score | Mean FC | P Value |
| --- | --- | --- | --- | --- |
| 1 | HDAC1 | 0.8333 | 1.7303 | <2.22E-16 |
| 2 | HDAC2 | 0.6667 | 1.5678 | <2.22E-16 |
| 3 | TTC38 | 0.5833 | 1.3717 | <2.22E-16 |
| 4 | HDAC10 | 0.4500 | 1.1170 | <2.22E-16 |
| 5 | PAH | 0.3500 | 1.1119 | <2.22E-16 |
| 7 | HDAC6 | 0.2833 | 1.1274 | 3.11E-12 |
| 700 | FADS2 | 0.0500 | 1.0069 | 0.2199 |
| 973 | Histone H4 | 0.0000 | 1.0347 | 0.9963 |

We further analysed a second 2D thermal profiling dataset interrogating ATP dose-dependent protein stabilisation in Jurkat crude lysate^14^. To assess the ability of TP-MAP to correctly identify direct targets of the metabolite ATP, we downloaded 4004 human proteins annotated with the gene ontology term “ATP Binding” (GO:0005524) from UniProt. Of these, 737 mapped to 6857 proteins (10.75%) proteins and annotated them as ATP-binding while the remainder were annotated as non-ATP-binding. Figure 3 depicts receiver operator characteristic (ROC) curves for detecting ATP-binding proteins using the TP-MAP stabilisation score at different threshold percentiles (ranging from 10% - 90%). The highest area-under-the-curve value was observed at 70% representing an upper cut-off of 1.39 and a lower cut-off of 0.73, with an AUC of 0.757. The mean fold-change may also be a viable predictor with an AUC of 0.746. As the study by Sridharan *et al.* provided data for additional replicates, we analysed all three available datasets with TP-MAP which gave comparable AUC values for replicates 2 and 3 (with an AUC of 0.758 and 0.737, respectively). There was a moderate correlation between mean fold-change and TP-MAP combined score (Spearman’s rank correlation, rho = 0.647, p < 0.0001), suggesting the two methods may be utilised in a complementary manner. At a p-value of < 0.0001, TP-MAP identifies 511 proteins as stabilised, of which 260 were annotated as stabilised at the 80% percentile threshold (50.88%) and 235 at the 70% percentile threshold (45.99%). In contrast, Sridharan *et al*. report identifying 753 proteins as stabilised at 1% FDR, of which 315 were annotated as ATP binding (41.83%). Among the top 753 proteins ranked using TP-MAP, we find 349 (46.35%) are annotated as ATP-binding, similar to those reported previously (Fisher’s exact test, p-value = 0.09).

**Figure 3.**
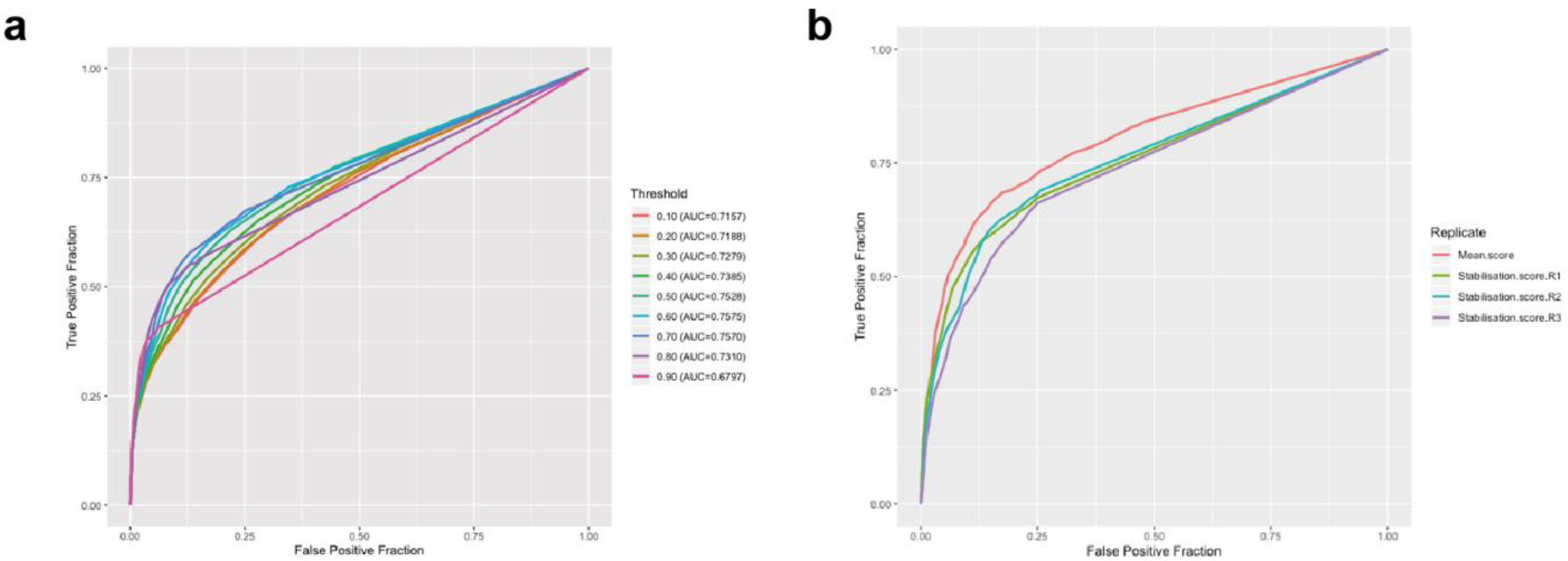
ROC curves classifying ATP-binding proteins using the TP-MAP 2D stabilisation score in 2D thermal profiling datasets using (a) varying TP-MAP thresholds and (b) in three replicates as well as the mean score between three replicates.

To investigate the capabilities of TP-MAP in exploring protein complex melting behaviour, we searched for proteasome subunits and identified PSMC4 as the most ATP-stabilised subunit. We then used the built-in CORUM functionality to identify known complexes that PSMC4 is a member of. The PA700 regulatory complex of the 26S proteasome included 19 additional proteins present within the dataset (Supplementary Figure 2). We applied the mean difference functionality in TP-MAP to identify proteins which exhibit similar aggregation behaviour and therefore might interact with PSMC4. The mean difference in melting behaviour between PA700 complex members and PSMC4 was 0.134 compared to 0.507 between PSMC4 and all other proteins (t-test, t = −13.9, df = 19.6, p-value < 0.0001). These results demonstrate that TP-MAP can aid in investigating protein complex co-aggregation^12^ and protein network mapping using built-in functionality.

## Discussion

Proteomics-based mechanism-of-action approaches are powerful means to elucidate the molecular interactions between drugs or metabolites in complex cellular systems. Many such studies have enabled the development of new therapeutics^23–25^ and these methods have now become an essential part of drug discovery programmes. Thermal profiling represents an attractive addition to the chemical biology toolbox as it does not require any modifications to the drug or compound of interest and works in intact living cells. However, in comparison to methods that are based on affinity enrichment before MS analysis, thermal profiling generates proteome-scale data for all conditions resulting in a significant amount of data. In the past, development of freely available and intuitive bioinformatics tools has supported the fast adaptation of new technologies in the systems biology community^26,27^. In order to facilitate data analysis, enable wet-lab scientists and therefore support further adoption of this technique, we developed TP-MAP as a data analysis tool for analysing thermal profiling datasets that provides an intuitive graphical user interface to conveniently explore large thermal profiling datasets and investigate proteins of interest via direct links to UniProt, STRING, and CORUM.

The main display is organised to allow a user to browse through a ranked list of hits and investigate the 1D melting curves or 2D matrices. In addition to the score, TP-MAP includes several other metrics classifying thermal shift data. For instance, 1D datasets include T_M_ and T_M_ shift for each replicate, RMSE for each replicate, and binary variables indicating whether a shift occurs in the same direction in both replicates and whether a shift between vehicle and treatment is greater than between the two vehicles. In 2D datasets, the mean fold-change is reported, as well as whether an effect is likely thermally induced or might be explained through changes in solubility or expression^14^ (i.e. a dose-dependent change is observed at the lowest temperature where a thermally induced shift would not be expected). This allows a user to identify hits and select proteins of interest for integrated downstream analysis, including investigating protein-protein interactions, gene-ontology functional enrichment, and identifying protein complex members (Supplementary Figure 2). Proteins of interest can be selected and exported as list in Excel file format whereas and curves can be stored as a PDF.

For 1D thermal profiling TP-MAP employs a scoring metric that incorporates curve fit quality metrics, including RMSE and reproducibility between replicates which compares well with recently proposed non-parametric methods that also incorporate curve-fit quality metrics^17^. This non-parametric method reportedly yielded 16 hits in a 1D TP dataset for panobinostat^5^ at a p-value threshold of 0.01, of which eight (HDAC1, −2, −6, −8, −10, H2AFV/H2AFZ, TTC38, ZFYVW28) are known panobinostat interactors. For comparison, the TP-MAP scoring identified the same eight proteins among the top 10 hits when a weighting was adjusted to 90%, and two additional known protein interactors (STX4 and HDAC7) were identified among the top 36 hits. The weighting option in TP-MAP is used to prioritise T_M_ shift over curve-fit quality. Depending on the target, we found that a weighting as low as 10-30% was optimal for identifying BTK as a dasatinib target, with a small but reproducible T_M_ shift (Figure 1), whereas optimal results for panobinostat were obtained at 90%, since targets exhibited large T_M_ shifts. Based on our global analysis of multiple 1D TP datasets, the default weight is set to 70%, however, TP-MAP includes functionality to change this setting interactively to explore hits using different weightings.

Importantly, the 1D TP-MAP score weights stabilised and destabilised proteins equally. Although the ATP dataset supports the notion that proteins become more stable when bound to a ligand, where a higher identification rate of ATP-binding proteins was obtained when only stabilised proteins were considered, our re-analysis of datasets identified known drug targets with negative shifts, such as BTK and YES1 for dasatinib, suggesting that negative shifts should be considered or else real targets could be missed, as has been supported by recent publications^28^. Notably, TP-MAP also identified the known downstream kinase effector STAT5B as an indirect target of dasatinib.

With TP-MAP we also introduce a new approach for ranking 2D thermal profiling datasets which does not rely on curve fitting. It has previously been noted that models that rely on fitting sigmoidal curves may not accurately represent the melting behaviour of all proteins in an organism, in particular proteins within specialised compartments, proteins within complexes, or proteins bound with a ligand^17,18^. TP-MAP allows a user to interactively change the cut-off at which a fold-change is considered stabilising or destabilising. This allows a user to limit scoring to proteins with the highest fold-changes. While at present TP-MAP does not have native support for multiple replicates in 2D TP datasets, we investigated whether predictive power could be improved by taking the average TP-MAP score across all three replicates in the 2D ATP dataset. We found the AUC of ROC curves to be higher when taking the average across three replicates, with replicates 1-3 having AUC values of 0.757, 0.758 and 0.737, respectively, whereas the mean score between the three replicates had an AUC of 0.802. We also considered an option to estimate the half-maximal effective concentration (EC50) from 2D thermal profiling datasets, although a reliable implementation has proved challenging as 2D TP datasets generally consist of few concentrations spanning several orders of magnitude. Further, we envisage the future integration of novel experimental methods based on thermal profiling such as proteome integral solubility alteration (PISA)^29^, into TP-MAP.

In summary, TP-MAP provides a GUI-based standalone application for analysing 1 and 2D thermal profiling datasets with added functionality to investigate targets of interest in the context of external databases without the need for deep *a priori* bioinformatics expertise. Therefore TP-MAP should facilitate further and broader adoption of thermal profiling-based approaches for target deconvolution and functional cellular mechanistic studies.

## Methods

### TP-MAP software

TP-MAP is a software application written in the Java programming language that has cross-platform support for Microsoft Windows, Apple macOS, and Linux operating systems. The software can import protein quantification files extracted from 1D and 2D thermal profiling experiments, and offers a graphical user interface to quickly identify proteins of interest, and provides additional functionality for exploring and analysing the data to determine likely targets identified in thermal profiling experiments. The software can import protein abundance levels either from TP-MAP formatted tables or ProteomeDiscoverer™ (Thermo Fisher Scientific, Waltham MA, USA) output in combination with an additional configuration table in which temperature and concentration values are specified for respective TMT samples and channels. An optional normalisation step can be performed whereby concentration-dependent ratios for 2D datasets are normalised by dividing by the median ratio at each temperature and concentration position.

### Scoring

#### 1D

TP-MAP supports thermal melting curve estimation for temperature dependent abundance ratios for vehicle and treatment, each consisting of two replicates. Up to four melting curves are fitted for vehicle and treatment replicates 1 and 2, and take the form of the following degradation curve equation^5^:

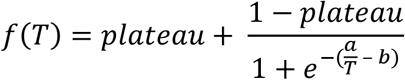

Where *T* refers to the temperature, and *a*, *b*, and *plateau* are constants.

For each protein, statistics about the curve fit are presented, including the melting point (T_M_) values for each vehicle and treatment condition (T_M_ V_1_, T_M_ V_2_, T_M_ V_3_ and T_M_ V_4_), thermal shift for each replicate (T_M_ shift V_1_T_1_ and T_M_ shift V_2_T_2_) as well as between vehicle conditions (T_M_ shift V_1_V_2_), root mean squared error for each fit (RMSE V_1_, RMSE V_2_, RMSE T_1_, RMSE T_2_), mean RMSE, as well as Boolean values indicating that both replicates shift in the same direction, and that the mean TM shift between vehicle and treatment is greater than between vehicle replicates (Delta VT > Delta VV). Finally, a score is calculated from these attributes to provide a suggested ranking.

Scoring of 1D melting curves is performed using a rank-based approach factoring in the Tm shifts of each replicate, RMSE of vehicle and treatment curve fits for each replicate, and mean difference between data points of each of the replicates, according to the following equation:

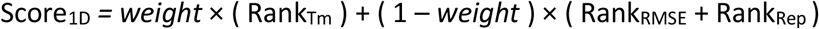

Where Rank_Tm_ is a vector (Tm_VT1_, Tm_VT2_) containing the scaled rank between [0,1] of increasing thermal melting point shift between vehicle and treatment in the two replicates respectively, Rank_RMSE_ is a vector (RMSE_V1_, RMSE_V2_, RMSE_T1_, RMSE_T2_) containing the scaled rank between [0,1] in decreasing order of root mean squared error (RMSE) of vehicle replicates 1 and 2, and treatment replicates 1 and 2, respectively, and Rank_Rep_ is a vector (Rep_V_ and Rep_T_) containing the scaled rank between [0,1] of the mean difference between vehicle replicates and treatment replicates, respectively. A *weight* is used to shift the score to prioritise either Tm shift or curve fit quality metrics. By default the *weight* is set to 70% and can be changed interactively, and depending on the quality of the dataset being analysed, may be shifted to prioritise curve fit quality or Tm shift. The resulting score is then scaled to range from 0 to 10.

#### 2D

A novel approach for scoring 2D heat maps is implemented, which is based on the assumption that concentration-dependent stabilisation or destabilisation will occur at an optimum temperature and concentration combination, and that adjacent temperature and concentration combinations will climb towards the optimum point. To achieve this, hill ascent and descent algorithms are implemented to identify peaks and troughs in a heatmap, respectively. TP-MAP calculates Score_stabilisation_ and Score_destabilisation_ ranging between 0 and 1, and Score_combined_ = Score_stabilisat–on_ - Score_destabilisation_ ranging from −1 to +1, where −1 indicates destabilisation and +1 indicates stabilisation. Supplementary Figure 3 depicts an example of how the scores are calculated. By default, a threshold beyond which a fold-change is considered stabilising or destabilising is set to the 80^th^ and 20^th^ percentile of the whole dataset and can be altered interactively by sliding the lower and upper threshold percentile sliders for destabilisation and stabilisation, respectively. A bootstrap analysis can optionally be run, whereby data points are resampled randomly for 10^6^ iterations to empirically estimate a P-value of the likelihood that *Score*_*2D*_ is reached by chance. In addition, the mean fold-change of proteins can be used as an additional measure to investigate proteins that show high rates of stabilisation or destabilisation and is displayed alongside *Score*_*2D*_, and may be used as alongside the TP-MAP score as a complementary method for identifying highly stabilised or destabilised proteins.

### Export format

TP-MAP allows a user to export selected proteins as Excel tables or as tab-separated text files. For 1D analysis, a PDF with melting curves can be exported.

### Downstream analysis of thermal profiling datasets

TP-MAP incorporates several methods for analysing 1D and 2D thermal profiling datasets. The table can be navigated interactively and proteins that appear to be of interest can be selected by ticking the checkbox in the first column.

The mean difference between a selected protein and all other proteins in the dataset can be calculated. The mean difference will be displayed in the “MD” column, which can be used to determine proteins that may share similar melting characteristics, such as protein co-aggregation^12^. Proteins with similar melting profiles can be identified by calculating the mean difference between a target protein of interest and the rest of the dataset. The mean difference is calculated from the difference in melting point ratio at each position in the heat map between two proteins and divided by the total number of positions.

In addition, analysing selected proteins with several external databases is possible. TP-MAP presently includes functionality for:

1. STRING^21^ network image: Load a protein-protein interaction network using the STRING database highlighting stabilised proteins as green and destabilised proteins as red (Supplementary Figure 2).
2. STRING functional enrichment: Load functional enrichment for selected proteins using the STRING database.
3. CORUM^30^: Identify protein complexes members of selected proteins using the comprehensive resource of mammalian protein complexes (CORUM) version 3.0, which includes 4274 mammalian protein complexes (Supplementary Figure 2).
4. UniProt^31^: Load UniProt pages for selected proteins.

### Implementation

The TP Meltome Analysis Program (TP-MAP) is implemented in Java 11, compiled using the Open Java Development Kit (OpenJDK) version 11.0.5 (https://adoptopenjdk.net). TP-MAP makes use of the following open source libraries: JavaFX version 11 (https://openjfx.io/), Apache Commons Math version 3.6.1 (https://commons.apache.org/proper/commons-math/), Apache HttpComponents version 4.5.10 (https://hc.apache.org/), JFreeChart version 1.6.0 and JFreeChartFX version 1.0.1 (http://www.jfree.org/jfreechart/), Apache PDFBox version 2.0.17 (https://pdfbox.apache.org/), and Apache POI version 4.1.0 (https://poi.apache.org/). TP-MAP can be run on computer operating systems with support for the Java Runtime Environment (JRE) version 11, including Windows, Mac OS X, and Linux.

## Acknowledgements

FF and KVMH are grateful for support by Myeloma UK and the Joyce and Norman Freed Trust. We would like to thank J. Ward and J. Stefaniak and all members of the Huber lab for testing the application and making insightful suggestions to improve the software.

## Availability of software

TP-MAP is licensed under a GNU GPL version 3 license (https://www.gnu.org/licenses/gpl-3.0.en.html). The source code and binary files are freely available for download at the following location: https://www.gitlab.com/ChemBioHub/tpmap

---

**Supplementary Figure 1.**
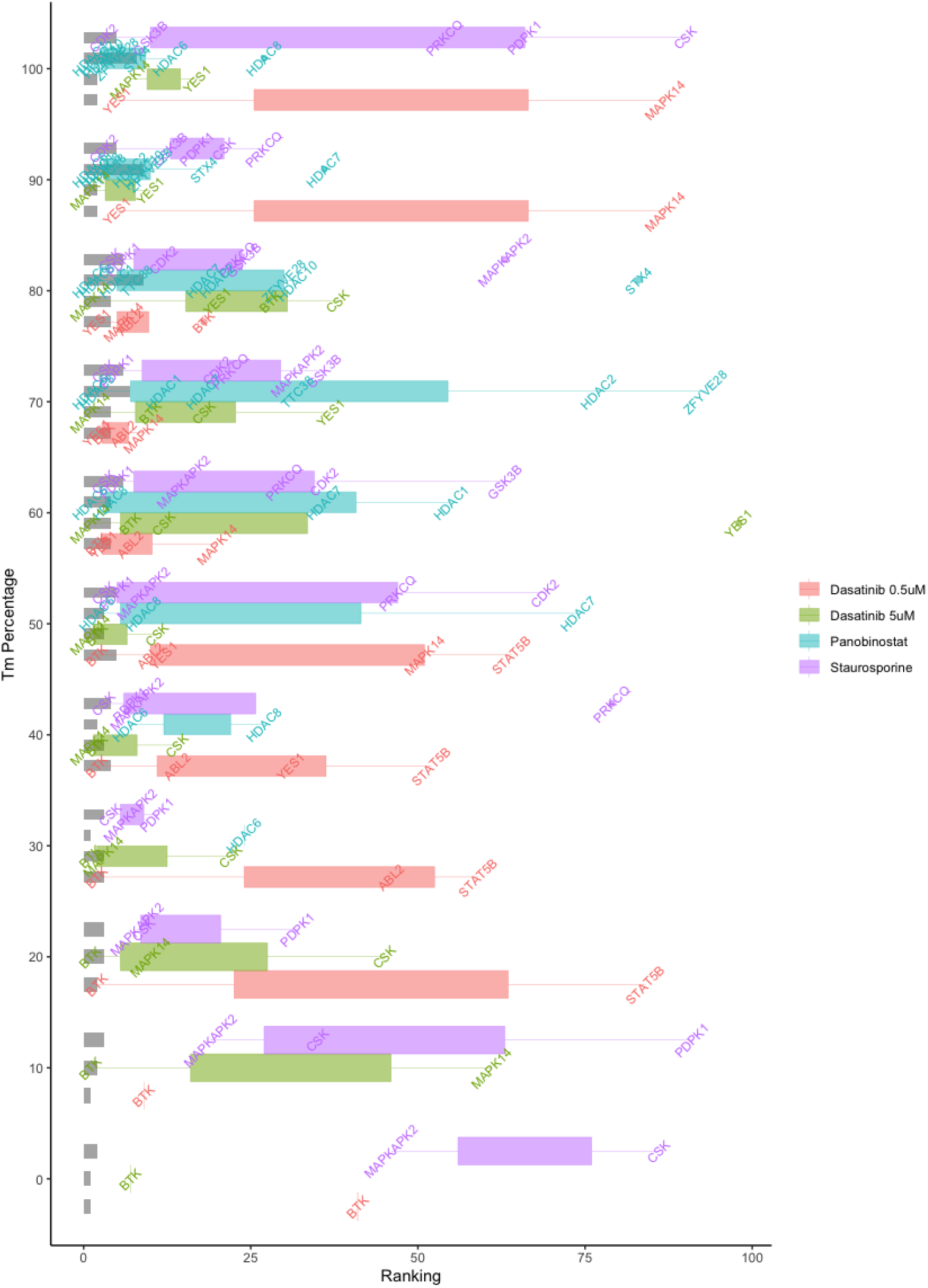
Ranking of targets in the top 100 within three 1D thermal profiling datasets when shifting the weighting of T_M_ shift between 0 and 100% at 10% intervals. Bars indicate the number of proteins identified for each drug within the top 100 hits, and box and whisker plots are used to show the distribution of rankings. Protein names are indicated at their respective rankings.

**Supplementary Figure 2.**
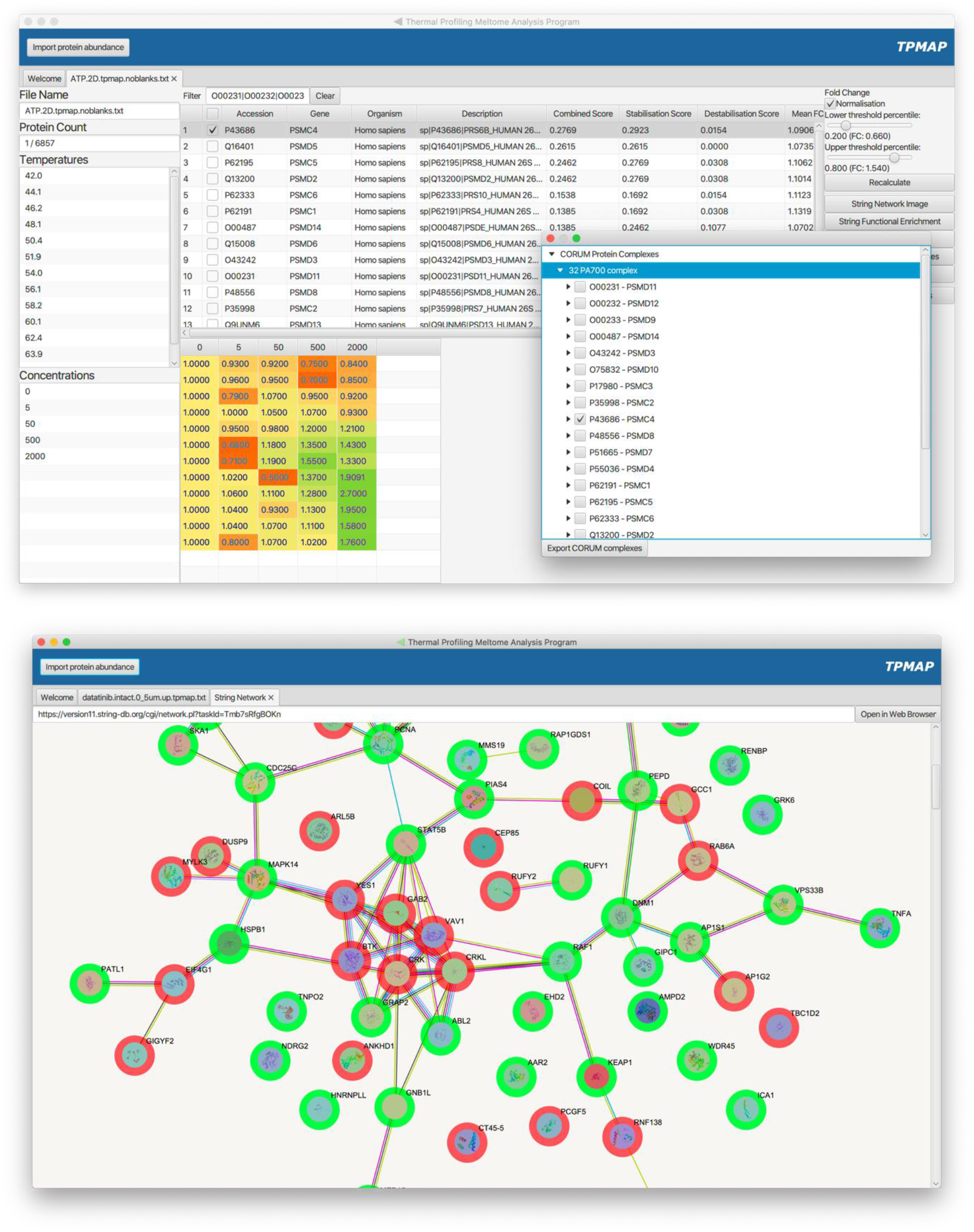
Screenshots displaying some of the downstream functionality of TP-MAP, including (a) identifying complex members of the proteasome in the ATP dataset and (b) a STRING network showing the top 75 hits for the dasatinib 1D data, showing a cluster of kinases and interacting proteins.

**Supplementary Figure 3.**
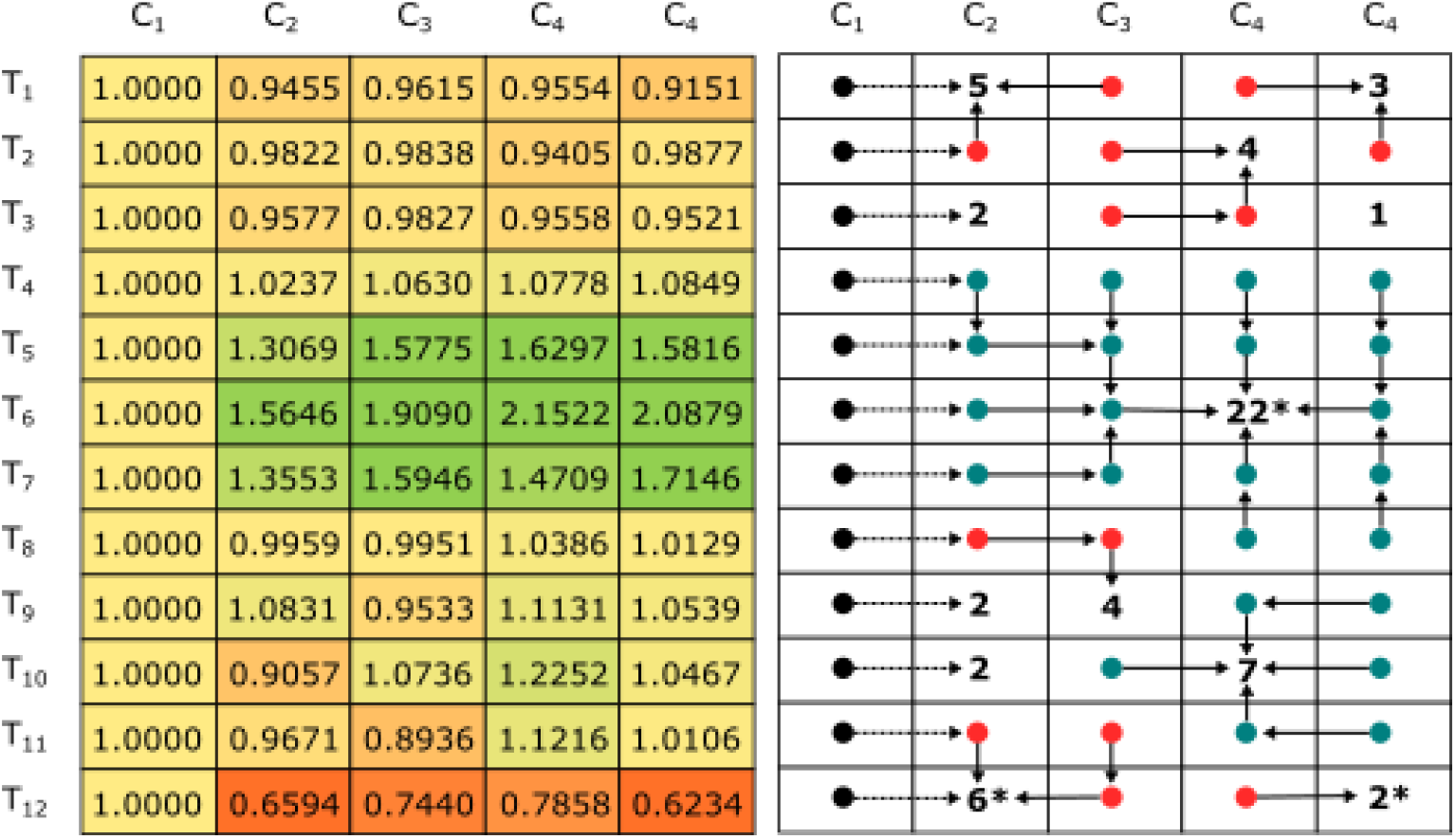
Schematic of the TP-MAP scoring algorithm, on the left a hypothetical concentration dependent stabilisation matrix is displayed with concentrations C_1_ to C_5_ and temperatures T_1_ to T_12_, on the right the corresponding TP-MAP scoring algorithm is shown. Teal and red nodes indicate stabilised (>1) and destabilised (<1) nodes climbing towards a peak or trough, respectively, which are represented by a number showing the number of nodes tending towards the respective peak or trough. Peaks and troughs marked with an asterisk (*) show those that are above or below a threshold of 1.5 or 0.75, respectively. Of these, Score_stabilisation_ is calculated as the number of nodes tending towards the highest peak (22 at T_6,_ C_4_) divided by the total number of nodes (60), 22/60 = 0.367, Score_destabilisation_ is analogously calculated as 6/60 = 0.100, and Score_combined_ is calculated as 0.367-0.100 = 0.267.

